# Machine learning model to predict risk assessment of a child inheriting a genetic disorder

**DOI:** 10.1101/2024.08.22.609095

**Authors:** Medha Ramaswamy, Shreya Senthilnathan, Adithiyan Rajan Indira Saravanan, M Snehal, Karthikeyan Siva

## Abstract

Advancements in Machine Learning (ML) have revolutionised precision medicine, particularly in predicting and preventing genetic disorders. This study provides a comprehensive analysis of ML models designed for the risk assessment of genetic disorders in children, based on clinical history and pedigree analysis. Leveraging a diverse dataset of familial genetic profiles, clinical outcomes, and medical histories, we employed ML algorithms to identify inheritance patterns, assess genetic risks for Mendelian disorders, and predict disease recurrence. The analysis incorporates several ML algorithms, including Gradient Boosting, XGBoost, Random Forest, Logistic Regression, Naive Bayes, and Support Vector Machines (SVM), with the Gradient Boosting model achieving the highest mean cross-validation score of over 0.99. Designed for primary care physicians and healthcare professionals, this model aids in genetic counselling by predicting genetic disorder recurrence based on family history. The paper also addresses ethical and legal considerations, emphasising the importance of genetic counselling and informed decision-making. This tool is intended to support, not replace, medical professionals. This work advances personalised risk assessment for Mendelian single-gene disorders, including autosomal dominant, autosomal recessive, and X-linked recessive disorders, contributing to the field of genomic medicine and facilitating effective family planning strategies.

**Author summary:** Genetic disorders are posing significant socio-economic, health, and psychological burdens due to the risk of inheritance, which can impact future generations with health complications. Genetic counselling has emerged as an important tool to help manage this issue, offering guidance to susceptible families. Recent advancements in Artificial Intelligence (AI) and Machine Learning (ML) have introduced new avenues for predicting the risk of genetic disorders with greater accuracy. The aim was to leverage machine learning models that would predict the risk assessment for single gene chromosomal Mendelian disorders. To train these models, patient data was collected from a clinical geneticist at a renowned hospital, all of whom had confirmed genetic testing results. This data was used to train the models that predicted the likelihood of the next child inheriting a genetic disorder. The initial results are promising, demonstrating a good fit with the data. However, it should be noted that larger sample sizes are needed to improve the accuracy of any model. With more extensive data, its predictive capabilities can be significantly enhanced. This tool has the potential to be a resource for genetic counsellors and primary healthcare physicians, aiding them in providing more accurate risk assessments and personalised guidance to families.

## Introduction

Genes and DNA are the basic building blocks of our heredity material, playing a fundamental role in shaping who we are and how we live our lives. Abnormalities or mutations in our genes cause genetic disorders, which disrupt the balance of our genetic makeup, leading to a wide array of health conditions and challenges. It is a condition where the normal DNA sequence is altered, either partially or completely. Genetic disorders affect more than 5% of live births and more than two-thirds of miscarriages [1,2]. Birth defects, syndromic diseases, chronic disabilities, neurological impairment, and other systemic illnesses are caused by single-gene disorders with Mendelian inheritance. Genetic disorders can result from the mutation of one gene (monogenic disorder) or mutations in multiple genes (multifactorial inheritance disorder or polygenic disorder) [3]. Diseases can be inherited from parents, or they can be caused by a spontaneous mutation in a specific gene. An inherited genetic disorder is present from birth, while a spontaneous mutation can occur either from birth or at a later point in life [4].

It is necessary to understand how genetic disorders are inherited from family members or de novo. A complete family medical history is a useful resource for demonstrating how diseases are inherited by subsequent generations. An individual has two copies of each gene, called an allele, one from either parent. Mutations that can occur on these genes affect how they work and can lead to a disease [5]. If the mutation leads to a disease, it is referred to as a pathogenic variation. Depending on the location of the gene and whether one or two normal copies of the gene are affected, diseases can be either autosomal dominant or autosomal recessive, respectively. Gregor Mendel initially noticed these patterns in garden pea plants, which is why this is frequently referred to as Mendelian inheritance [6].

There remains a notable lack of understanding of genetic disorders in India, even from healthcare experts. Raising awareness is very important for enabling early diagnosis, intervention, resulting in better patient outcomes and quality of life [3]. Public education on the importance of genetic testing, advocacy for genetic counselling services can reduce societal stigma and the impact of genetic disorders on individuals and families. This knowledge allows individuals to make informed choices about family planning and enhances the overall quality of healthcare services provided by professionals. It also helps in early diagnosis, intervention, treatment, and prevention of complications, which improves the quality of life of patients and their families [7].

With the existence of several genetic disorders, ranging from rare conditions like AA amyloidosis to more prevalent ones like thalassemia and cystic fibrosis, diagnostic technology is proving instrumental in increasing early detection and accurate diagnosis. Traditional methods of genetic risk assessment usually rely on family history and clinical evaluations commonly done by pedigree analysis, which may benefit from the integration of advanced computational approaches, especially when it comes to polygenic disorders and sporadic diseases [8]. Precise genetic diagnosis of single gene diseases using sequencing and other diagnostic methods aids in calculating the precise risk of recurrence.

Artificial intelligence (AI) is a general term that describes the use of computational algorithms capable of classifying, predicting, or drawing insightful conclusions from the analysis of large data sets. Machine learning (ML), a subset of AI, is the process of developing statistical models to predict outcomes or classify observations based on "training" supplied by humans using previously observed real-world data. These predictions are then applied to future data, while the new data is incorporated into the statistical model, which is constantly improving [9]. In recent years, machine learning has introduced innovative solutions to complex problems as it utilises large amounts of datasets and learns patterns and insights from them to make predictions, classifications, and decisions. ML has been proving to be very impactful in the healthcare field, helping with diagnosis, prediction, and treatment [10].

We have explored the use of ML in predicting the risk of getting a genetic disorder by training it with large sets of sample data. The primary objective is to enhance awareness and help in understanding the risk of recurrence in monogenic diseases, as well as to aid in the counselling efforts of primary physicians, medical specialists, and genetic counsellors. Additionally, it will support the evaluation of a couple’s risk for possible genetic disorders in their children, with a particular emphasis on Mendelian inheritance patterns. Initially, the focus will be on autosomal recessive, autosomal dominant, and X-linked diseases, utilising datasets comprising family history and genetic testing results from parents. The patients’ and, in some cases, the parents’ genetic test results were available. As comprehensive datasets become available, factors like demographics and regional differences will affect them. It can also provide valuable insights into public health trends. This can inform the development of policies, drive research on genetic disorders, and support genetic testing programs.

### Methods of Mendelian inheritance

**Table 1.** Types of Mendelian inheritance.

| Type | Affected parents | Risk of recurrence (in offspring) | Reference |
| --- | --- | --- | --- |
| Autosomal dominant | At least one affected parent or de novo | 50% if one parent is affected; less than 1% if de novo | [11, 12] |
| Autosomal recessive | Both parents are carriers, not necessarily affected | 25% in each pregnancy for the offspring |  |
| X-linked dominant | If the mother is affected, both males and females in the same generation may be affected. If the father is affected, all daughters and no sons will be affected | If mother is affected, 50% risk for females and males.<br>X linked diseases if severe, males may not survive or be lethal in utero to them if affected. |  |
| X-linked recessive | Males are generally affected and females | There is a 50% risk of males being |  |
|  | are carriers (since they carry only one X chromosome) but in cases with X inactivation females can be affected | affected. 50% risk of female being a carrier |  |

### Effect of consanguinity on inheritance

Consanguinity refers to a case where a couple shares a common ancestor (a common example would be when a couple are first cousins). Consanguinity is defined in degrees depending on the relationships: first degree, second degree, etc [13]. In many cultures, consanguinity is common. Consanguineous relationships in India are usually 2nd, 3rd, and 4th degree. The first degree of consanguinity is not possible. Children born to consanguineous parents are more likely to be at risk of autosomal recessive genetic diseases if they are carriers of a particular gene mutation with autosomal recessive pattern of inheritance. [14].

### Genetic disorders considered for ML training

The various Mendelian genetic disorders that were used for training and testing the ML model are given in Table 1.

**Table 2.**
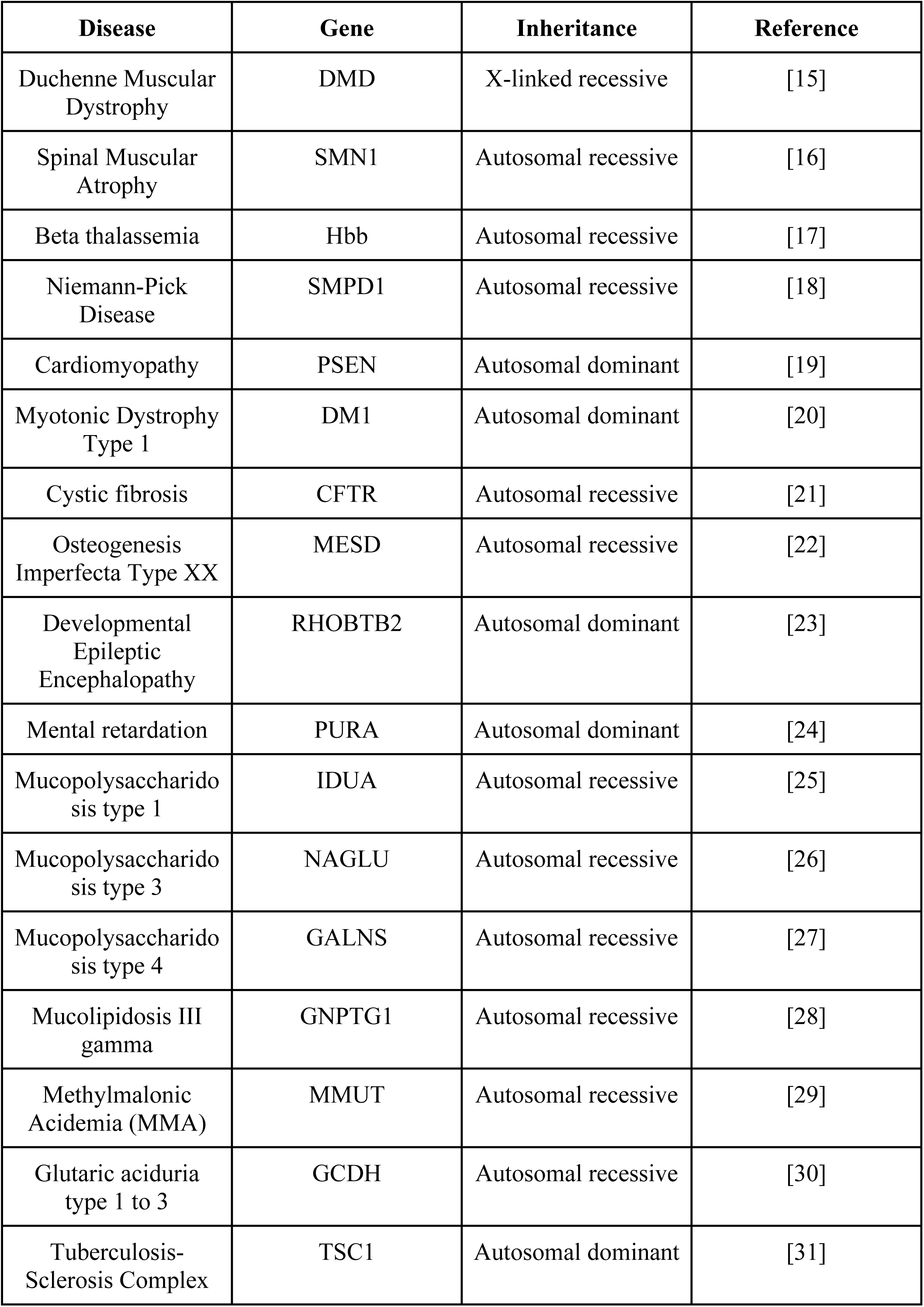
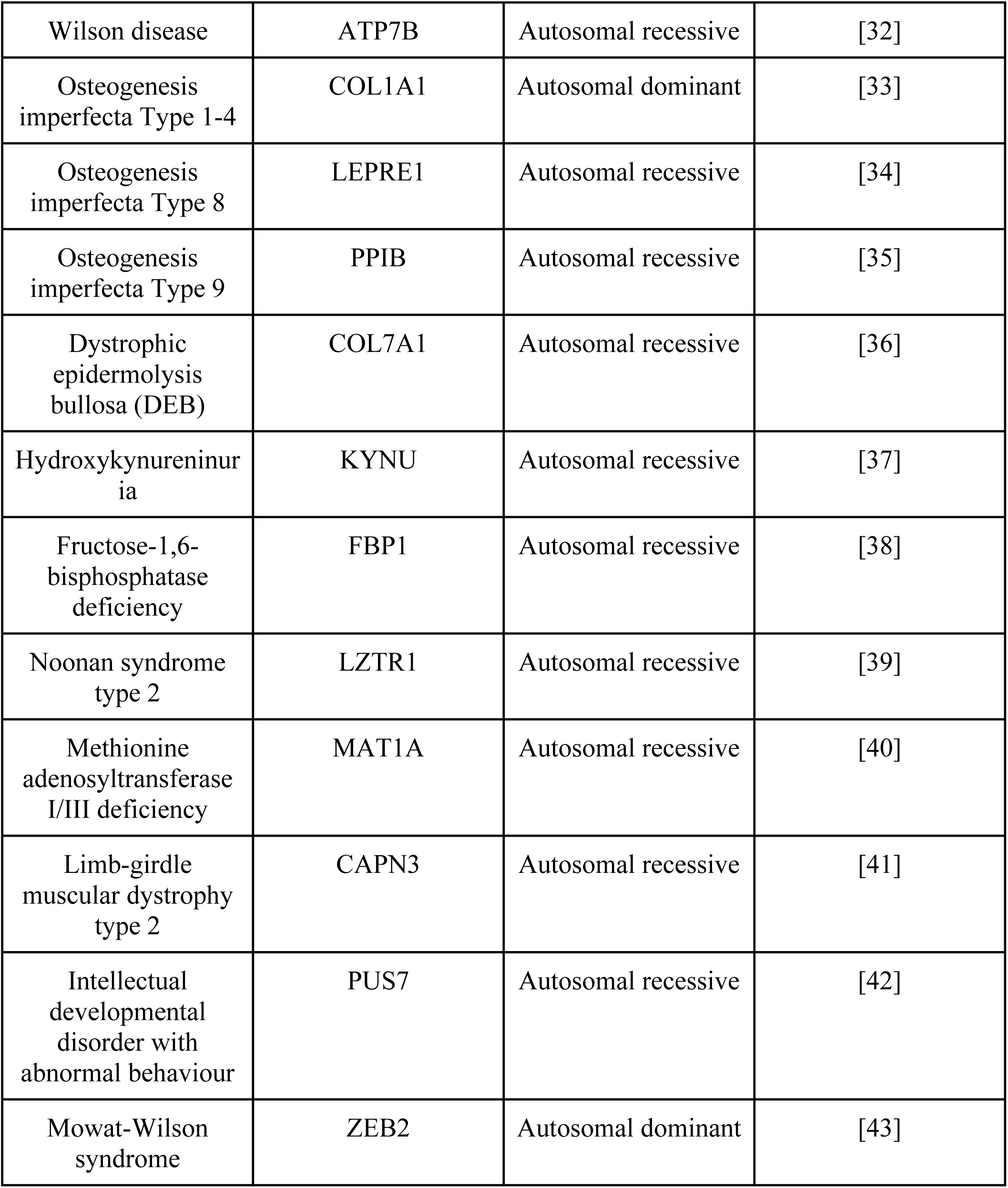
List of Mendelian inherited genetic disorders used in the dataset.

**Table 3.** List of columns used in the dataset.

| Column heading | Description |
| --- | --- |
| Type | Inheritance type |
| Genetic Disease | Name of the genetic disorder |
| Gene | Name of the gene responsible for disorder |
| Variation | Allele variation associated with disorder |
| Consanguinity | Indicates if the parents have a consanguineous relationship |
| Mother affected, Father affected | Indicates if the parent is affected by the disorder |
| Mother carrier, Father carrier | Indicates if the parent is a carrier of the disorder |
| 1st level | Indicates if any immediate family members are affected |
| 2nd level | Indicates if any extended family members are affected |
| G1, G2, G3, G4, G5 | The offspring are labelled from G1 to G5, with G1 being the first child and G5 being the fifth. Indicates if the offspring is affected |
| Risk assessment | Estimated probability of the offspring inheriting the disorder from the parents |

## Materials and methods

### Data collection

The patient data of 100 families with children affected by genetic disorders was collected from a genetic clinic, and a thorough clinical history was taken by a clinical geneticist and detailed examination, followed by genetic investigations and diagnosis through whole exome sequencing of the patient. Blood samples were processed in an accredited lab, and gene variations were classified according to American College of Medical Genetics and Genomics (ACMG) criteria for the dataset [44]. This included cases of DMD, SMA, beta thalassemia, Niemann-Pick, cardiomyopathy, myotonic dystrophy type 1, cystic fibrosis, osteogenesis imperfecta type XX, developmental epileptic encephalopathy, mental retardation, and many more. The columns in the dataset are “Type” (of inheritance), “Genetic Disease”, “Gene”, “Variation”, “Consanguinity”, “Mother affected”, “Mother Carrier”, “Father affected”, “Father Carrier”, “1st level family affected”, “2nd level family affected”, “G1”, “G2”, “G3”, “G4”, “G5”, and “Risk assessment." All cases have confirmed genetic testing and diagnosis.

### Data pre-processing

While preprocessing the data, a series of procedures were employed to ensure the quality of the dataset. Initially, irrelevant columns, such as identifiers and non-pertinent metadata, were removed to streamline the dataset for analysis.

Given their prevalence in genetic data, handling of missing values presented a challenge. Rather than using imputation methods, which could introduce inaccuracies, particularly in critical attributes, rows with missing values were dropped. This approach was necessary to maintain the accuracy and reliability of the dataset, minimising the risk of skewed results or erroneous conclusions. The target risk attribute was binned into three distinct categories to enhance predictive accuracy. This binning strategy simplified the prediction task and improved model performance, particularly with noisy or heterogeneous data.

Categorical variables were encoded using one-hot encoding techniques, transforming them into numerical representations suitable for machine learning algorithms. One-hot encoding was preferred for maintaining the structural coherence of categorical data while ensuring compatibility with various modelling approaches [45]. Due to the limited size of the dataset, which comprises approximately 130 rows, data augmentation techniques were employed to enhance model robustness and generalizability. Techniques such as resampling, bootstrapping, and SMOTE (Synthetic Minority Over-sampling Technique) were utilised. Resampling involved randomly selecting and duplicating instances from the original dataset, effectively increasing the sample size and improving performance stability. Bootstrapping creates multiple samples by random sampling with replacement, generating diverse training sets and reducing overfitting risks. This method provided a reliable estimate of the model’s predictive accuracy and variance [46]. SMOTE addressed class imbalance by generating synthetic samples for minority classes, balancing the class distribution, and improving predictive performance for underrepresented classes [47].

These data augmentation techniques expanded the dataset to approximately 1990 rows, significantly increasing the data available for training and evaluation. This augmentation addressed data scarcity concerns and contributed to the robustness and reliability of genetic disease risk assessment, ensuring more accurate and meaningful predictions across all classes.

### Data splitting

To further enhance reliability, the stratified k-fold cross-validation technique was employed. This method is crucial for medical datasets with class imbalances, as it ensures the preservation of class distribution in each fold, providing a representative sample of the overall dataset. The technique divides the dataset into k subsets (folds), where each fold serves as a validation set while the remaining k-1 folds are used for training, repeated k times. Stratified k-fold cross-validation allows for a comprehensive assessment of model performance by training and validating the model on different data subsets, thereby identifying potential overfitting or underfitting issues [48]. This technique maximises data utilisation, as each data point is used for both training and validation across different folds, enhancing the model’s generalizability and robustness. Performance metrics from each fold are averaged to provide a reliable estimate of the overall performance, ensuring that the evaluation is not biased by any single train-test split. By leveraging these data splitting techniques, the models achieved higher accuracy, generalizability, and robustness, offering valuable insights for genetic disorder prediction.

### Model selection and training

Various models were tested on this dataset to identify the most suitable one, including XGBoost, Gradient Boosting, Random Forest, Logistic Regression, Naive Bayes, and Support Vector Machines (SVM).

#### XGBoost

XGBoost (Extreme Gradient Boosting) is a scalable and highly effective implementation of the gradient boosting decision tree algorithm. It is an ensemble method that combines multiple weak decision tree models to create a strong predictive model. XGBoost employs a gradient-boosting framework that iteratively adds new trees to the ensemble, with each new tree correcting the errors of the previous ones. It includes advanced regularisation techniques and parallel processing capabilities, making it efficient for large-scale data analysis [49, 50].

#### Gradient Boosting

Gradient Boosting is an ensemble method that builds an additive model by sequentially training weak decision tree models, with each new model improving upon the errors of the previous models. It effectively handles complex and nonlinear relationships in the data by combining the predictions of multiple weak models to create a strong and accurate predictive model [51, 52].

#### Random Forest

Random Forest is an ensemble learning method that constructs multiple decision trees during training and outputs the class or prediction that is the mode of the classes or the mean prediction of the individual trees. It introduces randomness by training each tree on a random subset of the features and observations, reducing overfitting and improving generalisation. Random forests are known for their robustness, ability to handle high-dimensional data, and resistance to noise [53, 54].

#### Logistic Regression

Logistic regression is a statistical model that estimates the probability of a binary outcome based on one or more independent variables. It models the log-odds of the outcome as a linear combination of the predictor variables. Logistic regression is widely used for binary classification problems in machine learning [55, 56].

#### Naive Bayes

Naive Bayes is a probabilistic classifier based on Bayes’ theorem, which assumes independence among the features given the class label. It calculates the probability of each class given the feature values and predicts the class with the highest probability. Despite their simplicity and strong independence assumptions, Naive Bayes classifiers often perform well in practice, especially for high-dimensional data [57, 58].

#### Support Vector Machines (SVM)

SVMs are a class of supervised learning models that construct hyperplanes or decision boundaries in a high-dimensional space to separate different classes. SVMs aim to find the optimal hyperplane that maximises the margin between the classes, making them effective for classification tasks. Kernel functions are used to map the data into higher-dimensional spaces, allowing SVMs to model complex, non-linear relationships [59, 60].

## Results

All data and code used for model fitting and plotting is available on a GitHub repository at https://github.com/Adithiyan/genetic-disorder-risk-prediction.

The feature importance for the Logistic Regression model, presented in Figure 6, shows insights into the predictive factors of genetic disorder risk. It indicates the contribution of each feature/column to the model’s performance, measured using coefficients. These scores represent the influence of each feature on the model’s predictions, which can identify significant predictors. The plot displays the most important features, with the x-axis representing the feature importance scores and the features on y-axis.

**Figure 1.**
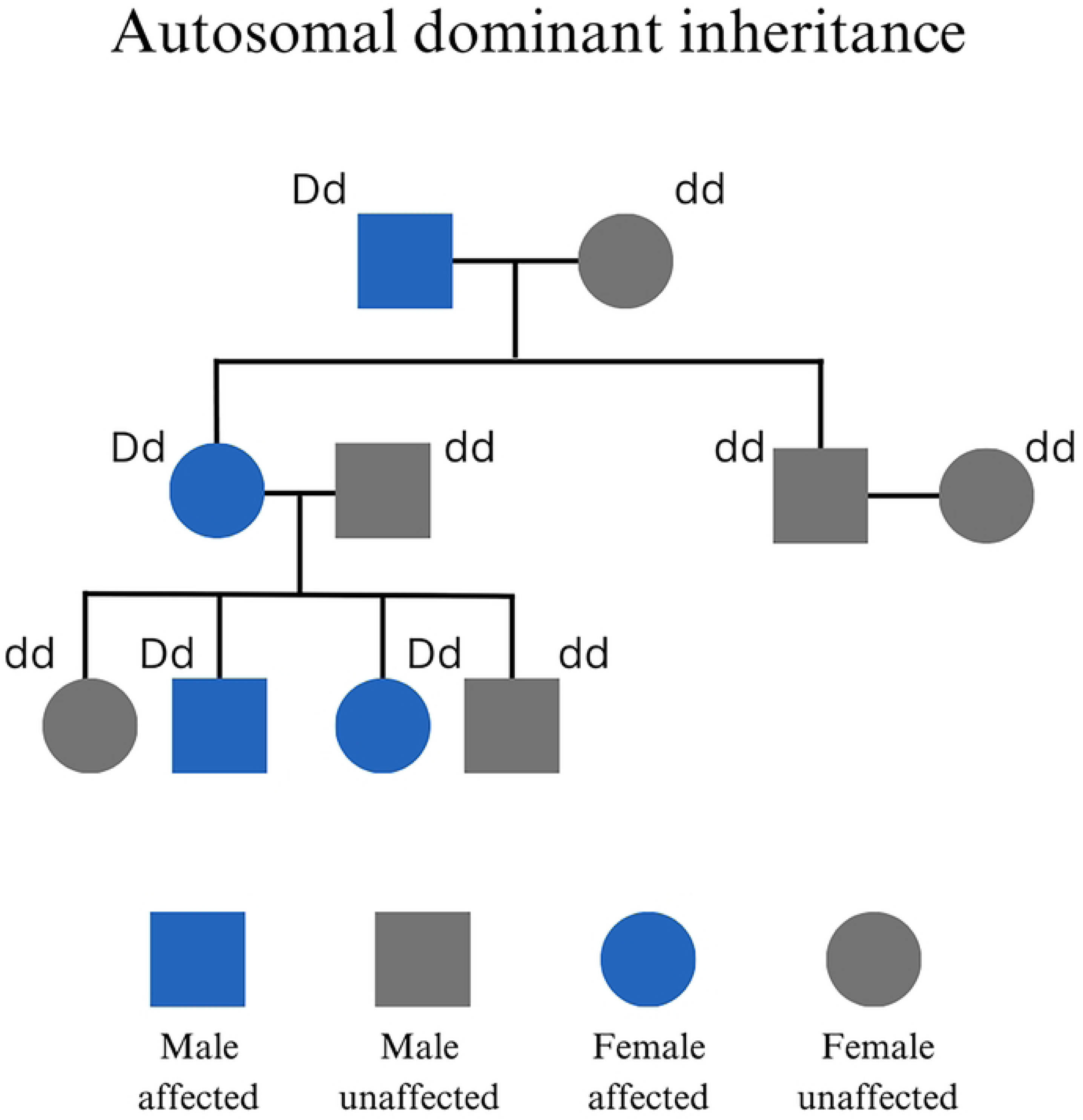
Pedigree charts depicting the inheritance pattern of the four major types of Mendelian inheritance.

**Figure 2.**
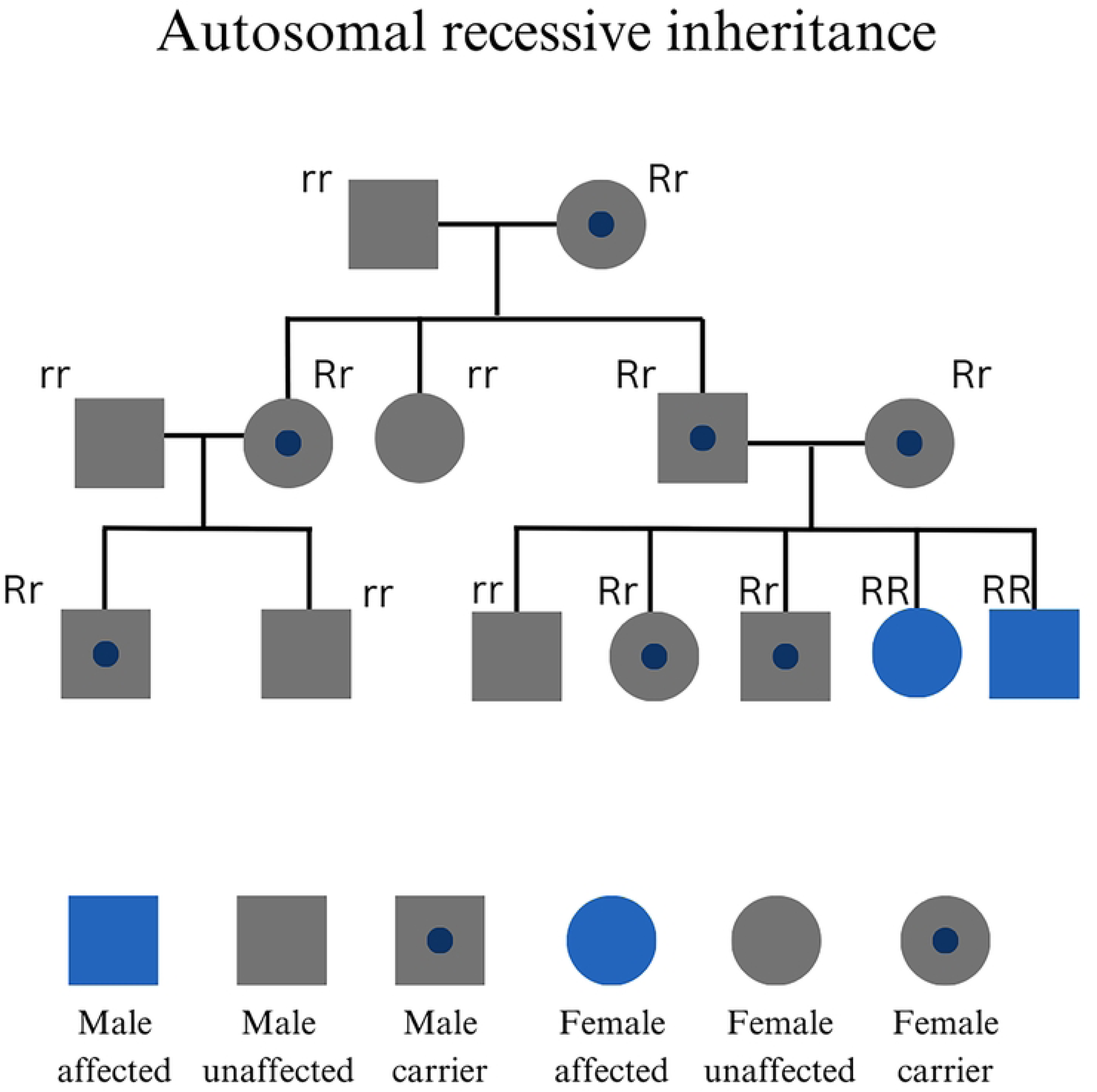
Pedigree charts depicting the inheritance pattern of the four major types of Mendelian inheritance.

**Figure 3.**
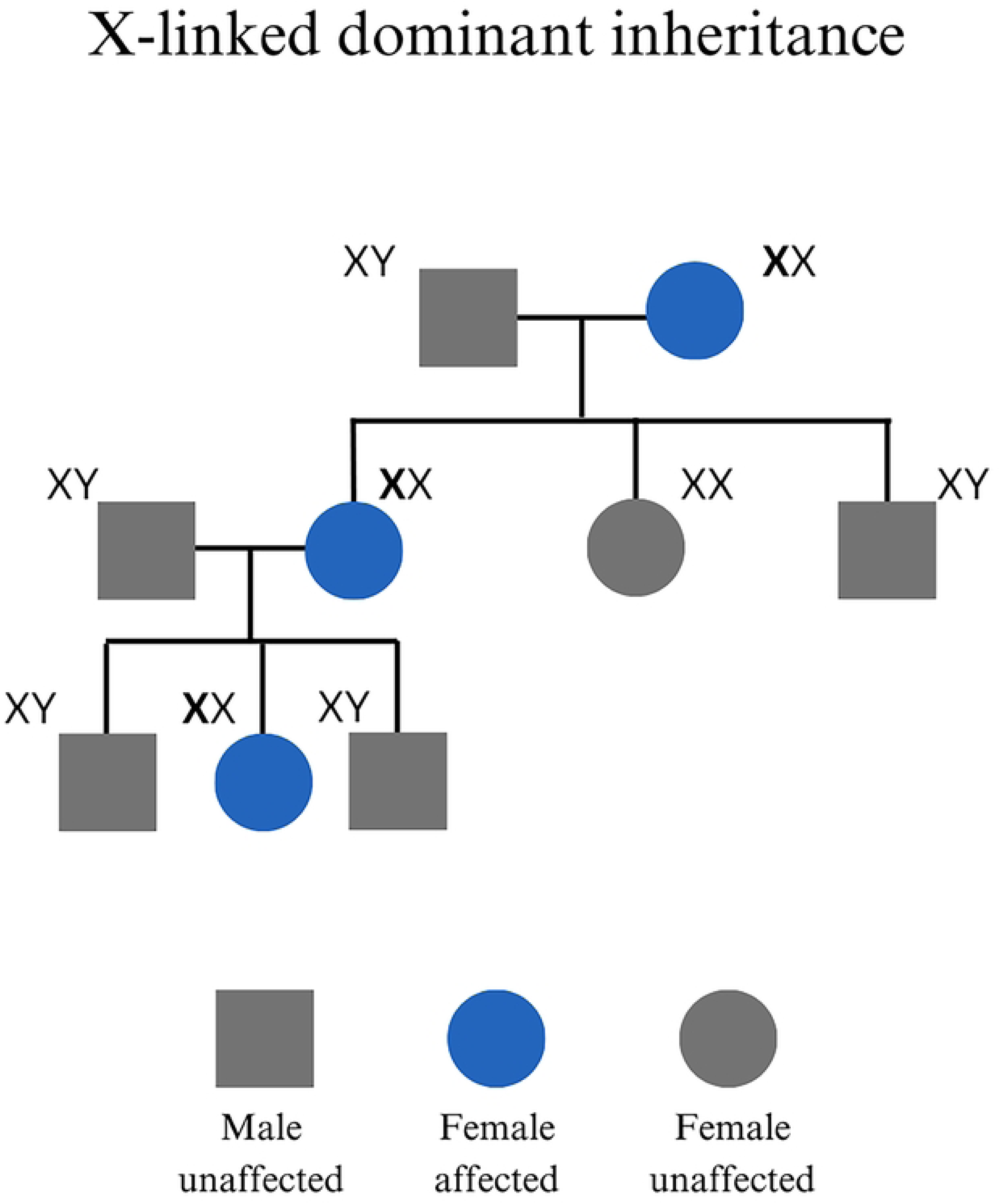
Pedigree charts depicting the inheritance pattern of the four major types of Mendelian inheritance.

**Figure 4.**
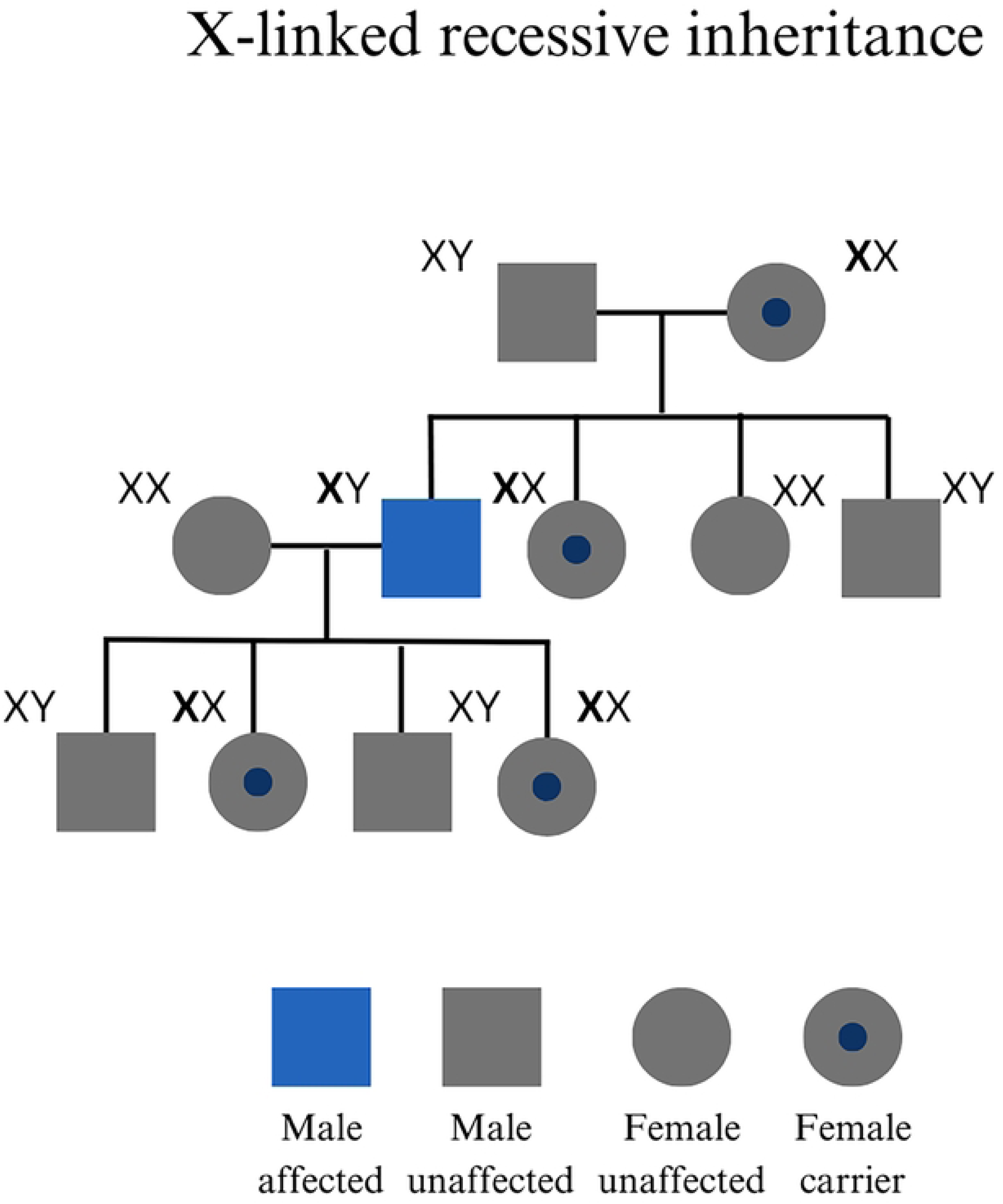
Pedigree charts depicting the inheritance pattern of the four major types of Mendelian inheritance.

**Figure 5.**
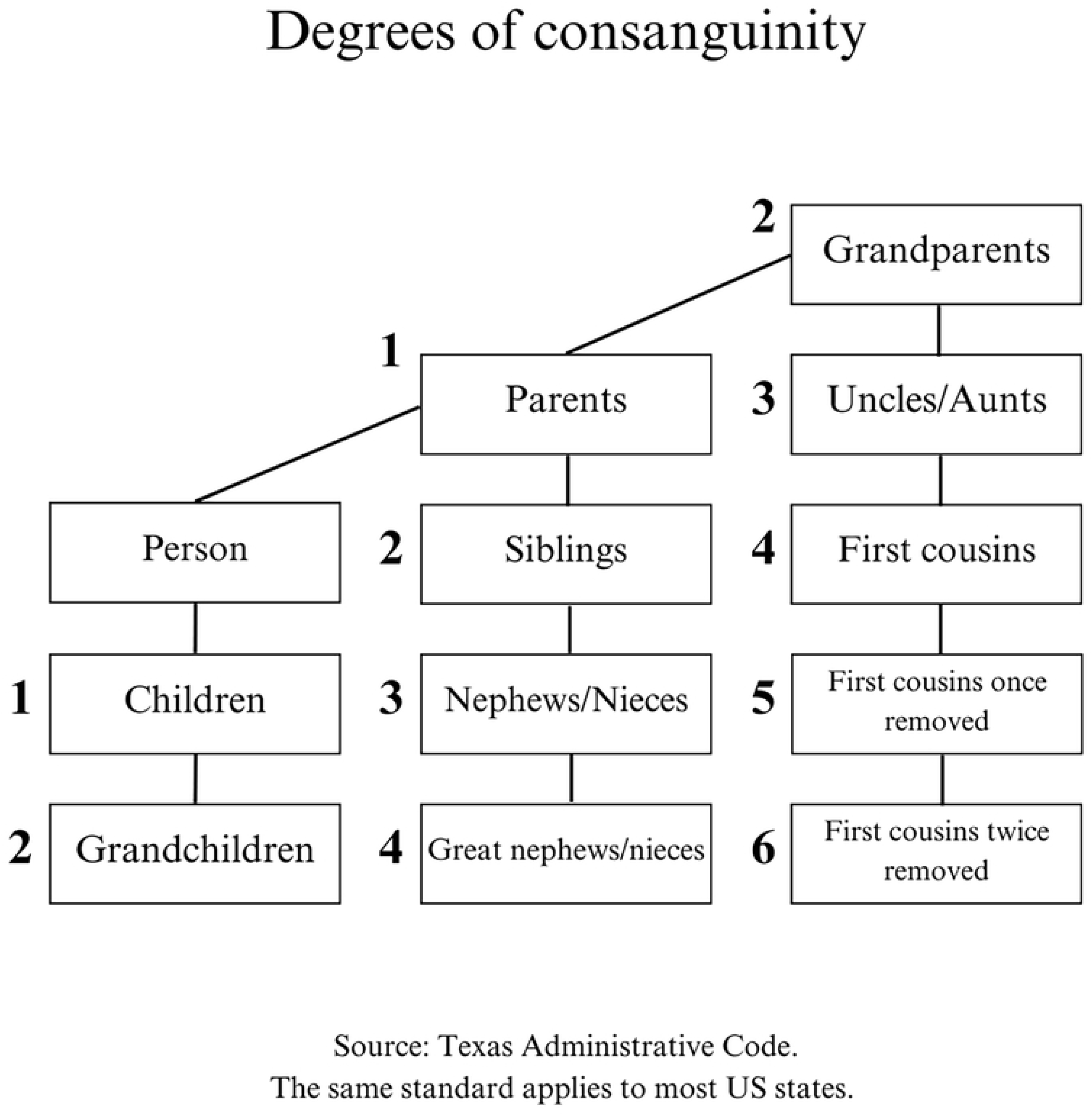
Degrees of consanguinity with relation to an individual.

**Figure 6.**
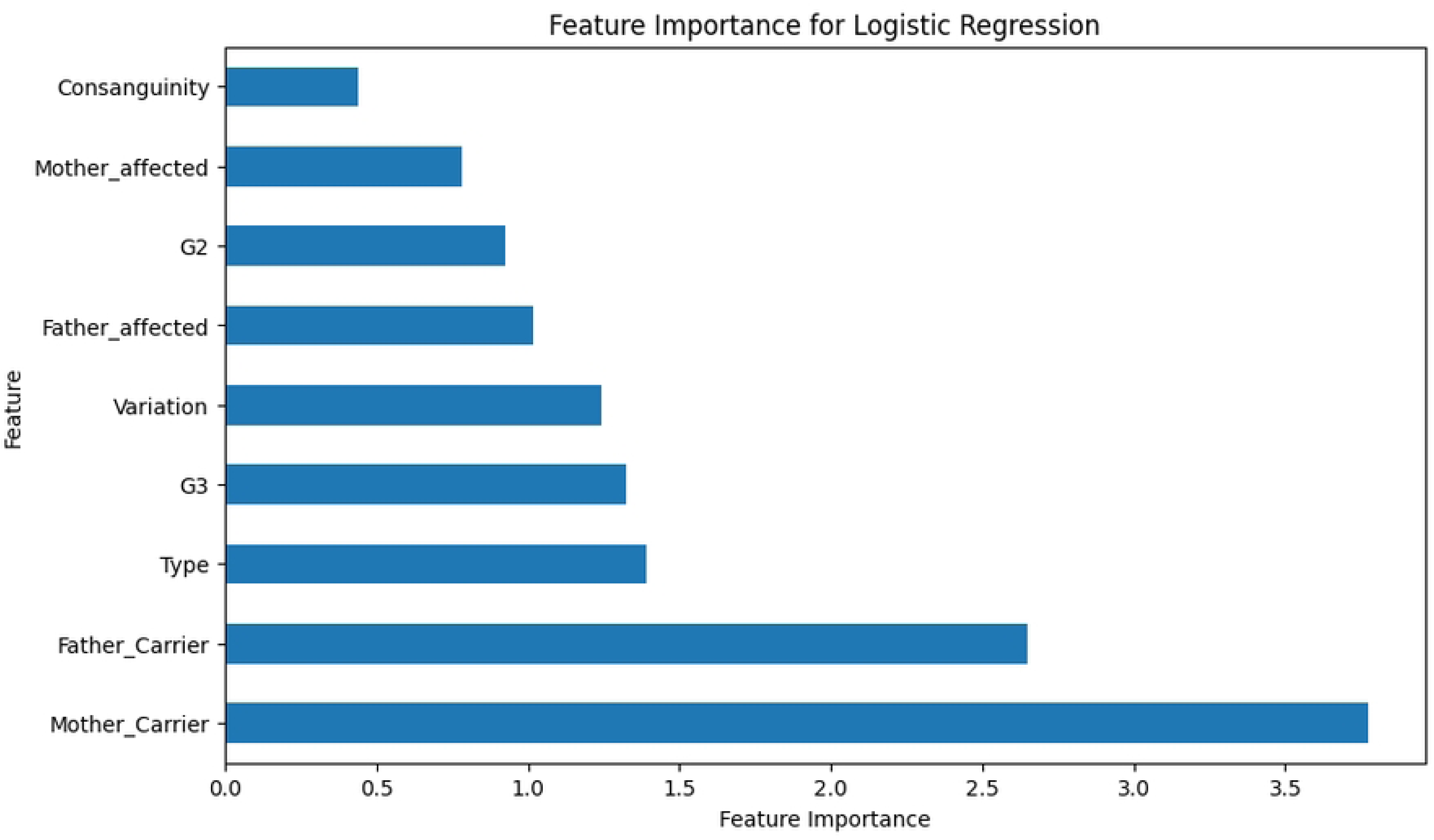
Feature importance for Logistic Regression.

‘Mother_Carrier’ exhibited the highest feature importance score, indicating that the mother’s carrier status plays an important role in determining genetic disorder risk. ‘Father_Carrier’ closely follows, highlighting the father’s carrier status’s significant contribution to risk prediction. However, this finding might be attributed to a potential skew in the dataset towards the mother’s features, with the mother being a carrier in more cases. Other features with high importance scores include ’Type’, denoting the mode of genetic inheritance; ’Variation’, indicating the zygosity of the gene; and G2 and G3, reflecting the affected status of the second and third offspring, respectively; the affected status of the father and mother; and consanguinity, all of which provide valuable information for risk prediction.

Features related to the parents’ carrier status and genetic factors appear to be more influential than features such as consanguinity in predicting the risk of genetic disorders. This insight emphasises the significant role of parental genetic status in predictive models and can guide future research and risk assessment strategies.

The performance of various machine learning models was evaluated for predicting the risk of a genetic disorder using cross-validation. The mean cross-validation scores for each model are presented in Figure 7. This metric reflects the average performance of the model across multiple iterations of cross-validation, providing an estimate of its generalisation ability and accuracy on unseen data. A higher mean cross-validation score indicates superior predictive performance, highlighting the model’s efficacy in accurately assessing genetic disorder risk in clinical settings.

**Figure 7.**
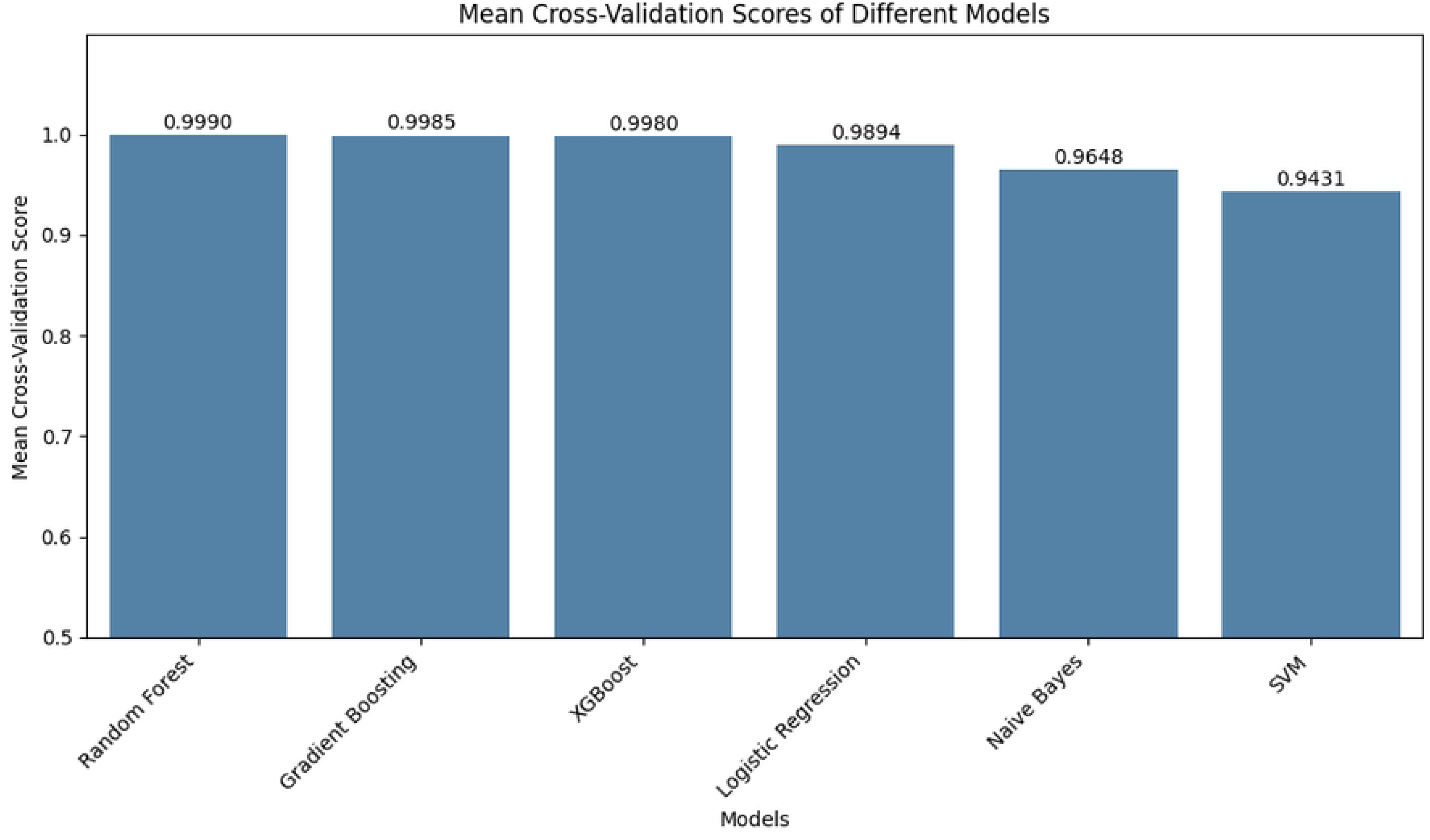
Mean cross-validation scores of different models.

The Gradient Boosting model achieved the highest mean cross-validation score of 0.9990, indicating excellent predictive performance. The XGBoost and Random Forest models, as well as decision tree-based ensemble methods, demonstrated strong predictive capabilities with mean cross-validation scores of 0.9980 and 0.9975, respectively. The Logistic Regression model achieved a mean cross-validation score of 0.9843, representing reasonably good predictive ability. The Naive Bayes model obtained a score of 0.9611, slightly lower than the other models. The Support Vector Machine (SVM) model exhibited the lowest mean cross-validation score of 0.8640 among the evaluated models. It is noteworthy that the XGBoost, Gradient Boosting, and Random Forest models, which achieved the highest scores, are decision tree-based ensemble methods that combine multiple decision trees to improve performance and reduce overfitting. However, the exceptionally high cross-validation scores, particularly for XGBoost, raise concerns about potential overfitting, where the models may have captured specific patterns or idiosyncrasies in the data that may not generalise well to unseen data. Overall, the results suggest promising predictive capabilities across all models, with decision tree-based ensemble methods outperforming others.

## Discussion

### Ethical issues

These machine learning models are designed to support, not replace, clinical diagnosis. Individuals should always consult registered medical practitioners before making any clinical decisions related to family planning or health. The models assist primary consultants in diagnosing and counselling patients and their families about the risk of disorder recurrence, complementing rather than replacing clinical expertise. Patient data used for training is fully anonymized to maintain confidentiality. These models are intended exclusively for the risk assessment of Mendelian disorders and are not to be used for a definitive diagnosis.

Confirmatory diagnoses must be made by registered medical practitioners to ensure accurate disease management and informed family counselling.

### Limitations

It should be acknowledged that this model has certain limitations. It is trained only on Mendelian inheritance patterns for monogenic disorders, excluding polygenic diseases involving multiple genes. Therefore, the model’s outputs are valid exclusively for monogenic disease inheritance risks, including autosomal dominant, autosomal recessive, X-linked dominant, and X-linked recessive conditions. Additionally, the relatively small sample size of the training data restricts predictive accuracy and generalisability. To improve the model’s ability to accurately access and predict genetic disorder risks across different populations, it is important to expand the training dataset to include a more diverse range of genetic profiles. This will also enhance the model’s robustness and precision.

### Future prospects

The capabilities of this model will be expanded to include polygenic disorders and take into account lifestyle factors. Its application can be extended to digenic and mitochondrial conditions by increasing the sample size through additional clinical data collections. This also ensures that more diverse populations are represented within the dataset. Artificial intelligence and machine learning play crucial roles in healthcare research, especially with regard to managing large volumes of patient data [6]. Our aim is to maintain data integrity, ensure privacy, and implement robust security measures while integrating patient data from various sources. Hosting the model locally will enhance both data security and patient confidentiality.

Ethical considerations, including equitable access and prevention of genetic information misuse, will remain central throughout development.

Enhancing the model will not only improve genetic risk assessments but can also elevate the genetic counselling proficiency of primary healthcare workers, significantly impacting patient outcomes. Rigorous validation in diverse clinical settings will be pivotal in supporting the model’s reliability and effectiveness. Continual refinement and expansion will also play a critical role in advancing precision medicine and enhancing healthcare outcomes.

## Conclusion

Machine learning’s application in genetic disorder risk assessment is a major step forward in healthcare, simplifying disease analysis for primary healthcare. This paper details the methodology for creating and evaluating models, focusing on Mendelian inheritance patterns using diverse datasets of clinical information.

The results show promising predictive abilities across various machine learning models, especially decision tree-based ensembles. However, it is crucial to address potential overfitting, as well as to validate the models for robustness, generalisability, and security. Techniques like regularisation or feature selection can help enhance generalisation. These models significantly improve personalised risk assessment for genetic disorders, aiding informed decision-making in family planning. It can help in enhancing the quality of counselling by primary healthcare providers.

Expanding this framework allows clinicians and researchers to use its capabilities more effectively, leading to better family planning. The comprehensive risk assessment model presented here is a useful resource for clinicians and genetic counsellors and is expected to be widely adopted in clinical practice after validation with larger and more diverse populations. This advancement will allow for more informed decision-making, thus benefiting individuals and families affected by genetic conditions as genomic medicine progresses with technology.

